# PRL1 negatively regulates Rho GTPase-independent and - dependent signaling pathways maintaining actin microfilament dynamic for pavement cell morphogenesis

**DOI:** 10.1101/2023.02.28.530536

**Authors:** Xiaowei Gao, Bo Yang, Jingjing Zhang, Chi Wang, Huibo Ren, Ying Fu, Zhenbiao Yang

## Abstract

Actin dynamic is critical for cell morphogenesis in plants, but the signaling mechanisms underlying its regulation are not well understood. Here we found *PRL1* (<u>P</u>leiotropic <u>R</u>egulatory <u>L</u>ocus1) modulates leaf pavement cell (PC) morphogenesis in Arabidopsis by maintaining the dynamic homeostasis of actin microfilaments (MF). Our previous studies indicated PC shape formation was mediated by the counteracting ROP2 and ROP6 signaling pathways that promote the organization of cortical MF and microtubules (MT), respectively. Our genetic screen for *ROP6* enhancers identified *prl1* alleles. Genetic analysis suggested that *prl1* acted synergistically with *ROP2* and *ROP6* in regulation of PC morphogenesis. We further found that the activities of ROP2 and ROP6 were increased and decreased in *prl1* mutants, respectively. Interestingly *prl1* was found to prefer to depolymerize MF independent of ROP2 and ROP6. Stress (high salinity and low temperature) induced similar changes of ROP activities as do *prl1* mutations. Together our findings provided evidence that PRL1 governed two signaling pathways that counteractively maintain actin dynamics and resultant cell morphogenesis.

## Introduction

Cortical actin microfilaments (MF) play pivotal roles in a wide variety of cellular processes (Chhabra and Higgs, 2007). In particular they are critical for cell polarization, polar growth, morphogenesis, and movement in eukaryotic cells (Li and Gundersen, 2008). In plants, cortical MFs have been implicated in tip growth, cell morphogenesis, cell polarity establishment, and defense (Hussey et al., 2006; Li et al., 2018 (Chang et al., 2017; Fu et al., 2005; Fu et al., 2002; Huang et al., 2015; Li and Staiger, 2018; Szymanski and Staiger, 2018)). The specific roles of cortical AF rely on the existence of their diverse forms as well as their dynamics (Campellone and Welch, 2010; Chhabra and Higgs, 2007). The organization of specific AF and dynamics are mediated by a series of actin-binding proteins (ABPs) such as actin nucleation factors, actin cross-linking proteins, capping proteins, G-actin sequestering proteins, actin depolymerizing factors (ADFs) (Campellone and Welch, 2010; Edwards et al., 2014; Kudryashova et al., 2017; Pollard, 2007; Van Troys et al., 2008).

The activity of these ABPs is regulated by intracellular and extracellular signals. In plants signaling mechanisms modulating ABPs are emerging but remained poorly understood. Plant-specific ROP (Rho-like GTPase from plants) subfamily of the conserved Rho family of small GTPases, well known conserved molecular switch that control actin organization and dynamic in eukaryotes, were among the first signaling proteins known to regulate actin organization and dynamics in plants (Smith and Oppenheimer, 2005; Yang, 2008; Fu et al., 2005; Fu et al., 2002; Fu et al., 2009; Humphries et al., 2011). In Arabidopsis, auxin promotes formation of the puzzle shape of leaf pavement cells by activating the antagonistic ROP2/ROP4 and ROP6 signaling pathways (Fu et al., 2005; Xu 2010). ROP2 and ROP4 are functionally redundant in promoting lobe formation via their promotion of cortical fine AF accumulation, and ROP6 promotes indentation by organizing ordered cortical MTs in the indentation regions. Other hormones such as brassinosteroids and cytokinin and environment factors also regulate pavement cell morphogenesis (Higaki et al., 2020; Li et al., 2013; Li et al., 2020; Liu et al., 2018), but mechanisms by which these internal and external signals regulate this process remain unclear.

*PRL1* (**P**leiotropic **R**egulatory **L**ocus 1) was initially isolated in a null mutant exhibiting multiple development defects including inhibited elongation of primary roots and hypocotyls and hypersensitive development responses to a variety of signals in Arabidopsis (Nemeth et al., 1998). Additional role of PRL1 in plant defense was discovered in company with ATCDC5, MOS4, MAC3A and MAC3B in MAC/Prp19/NTC complex (Monaghan et al., 2009; Palma et al., 2007), but functional mechanisms are unknown. It was also uncovered that PRL1 was a negative regulator for SnRK1 in E3 ligase complex in the regulation of stress or intracellular energy deprivation response (Abraham et al., 2003; Baruah et al., 2009; Salchert et al., 1998). PRL1 was also reported to influence miRNA production and alternative splicing as a regulatory component interacting with core components like DCL1, DCL3/4 and HYL1 (Jia et al., 2017; Li et al., 2018; Zhang et al., 2014). In yeast system, it was revealed that *PRL1* was involved in regulation of actin configuration and polar cell development (Xia et al., 1996), but no followup work has been conducted to explore such specific role of *PRL1* in plants.

Here, we identified PRL1 as a new regulator of pavement cell morphogenesis via its role in the maintenance of actin assembly and ROP signaling-correlated actin organization. Genetic analysis confirmed *ROP2/6* and *prl1* interacted synergistically in shaping PC. Further cytological and biochemical analysis revealed that *prl1* caused depolymeriztion of cortical microfilament and changes of ROP2 and ROP6 activities. Thus, our results revealed that PRL1 was a key regulator of dynamic actin microfilament homeostasis that is essential for cell morphogenesis in Arabidopsis.

## Results

### The *prl1* mutation caused aberrant cotyledon pavement cell shape in *Arabidopsis*

To identify new components in the ROP signaling pathways that regulate pavement cell morphogenesis, we generated a mutant population by EMS–induced mutagenesis in a line that moderately over-expressed *ROP6, ROP6-5* (*GFP-ROP6*) (Fu et al., 2009). Compared to WT, *ROP6-5* line possesses slightly narrower necks in leaf pavement cell and almost identical lobe density (Figure 1A, 1B, 1K and 1L). A recessive mutant *L171* (*prl1-10 ROP6-5*) was isolated, in which the contour of pavement cells was affected severely with the jigsaw-puzzle pattern nearly completely lost at 4^th^ day after stratification (Figure 1A and 1D). The quantification showed that the average lobe density of *L171* was decreased to 0.12±0.03/1000μm^2^, compared with those of WT (1.70±0.17/1000μm^2^) and *ROP6-5*(1.73±0.10/1000μm^2^) (t-test, p<0.001); the mean neck width of *L171* pavement cell was 23.75μm, which increased significantly when compared to those of WT (17.83±0.29μm) and *ROP6-5* (16.77±0.68μm) (t-test, p<0.001) (Figure 1K, 1L). The increase in neck width of pavement cells suggests that the cell outgrowth was promoted in *prl1* mutant and the reduction of lobe density suggests that cell polarity became reduced. In addition, we found that the length of primary roots in *L171* was dramatically reduced (Figure 1I), a property genetically confirmed to be co-segregated with pavement cell phenotype and helpful for us to efficiently identify homozygotes in the F2 population for the map-based cloning.

**Figure 1.**
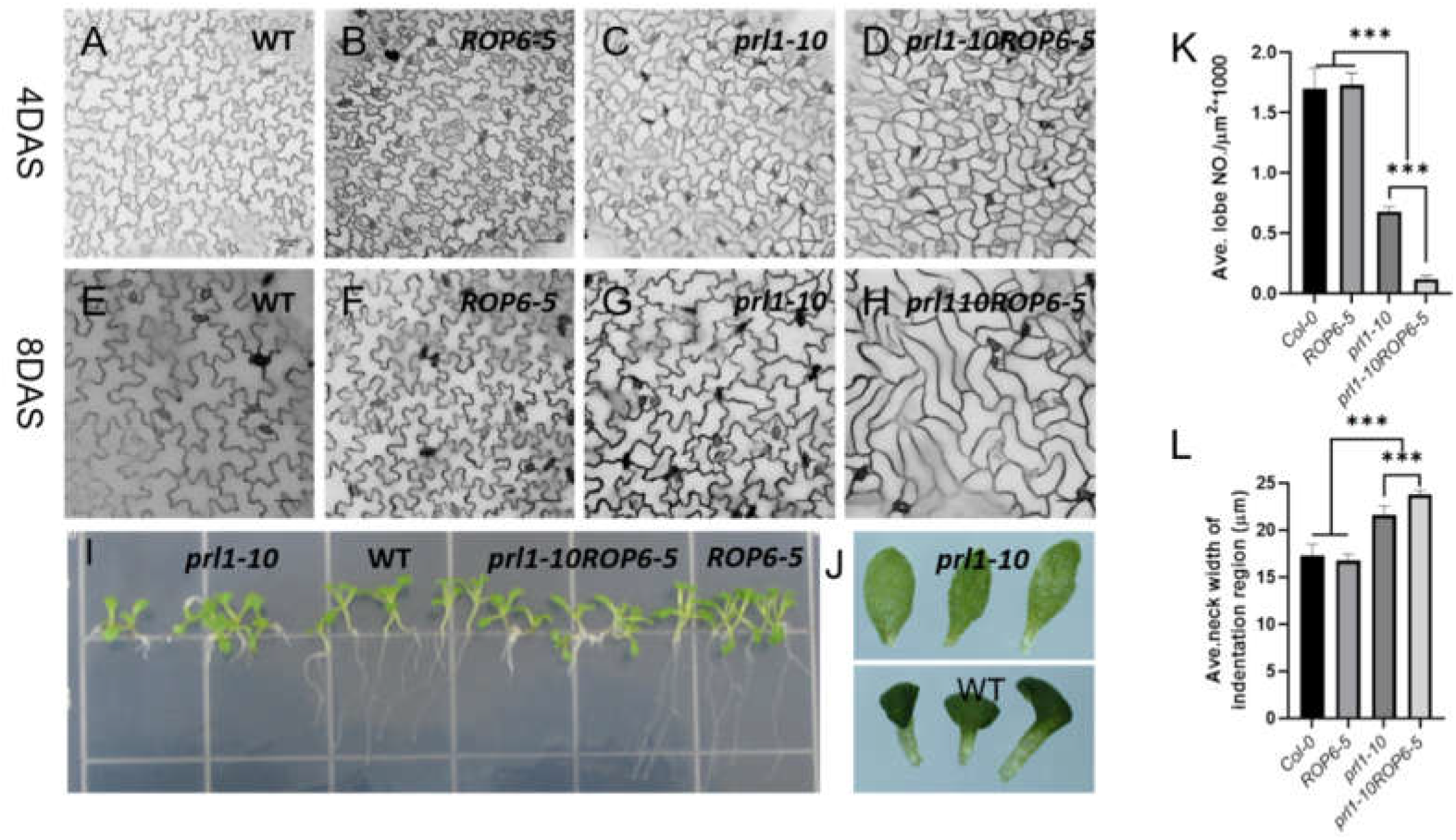
The *PRL1* mutation dramatically alters PC polarity and augments effect of ROP6 on polar PC development. (**A-D**) the cotyledon pavement cell contours of Col-0, *ROP6-5, prl1-10* and *prl1-10ROP6-5* at 4^th^ day after stratification (DAS), mutation in PRL1 dramatically reduced interdigitation of pavement cell in cotyledon. Scale bar, 50μm. (**E-H**) The cotyledon pavement cell contours of Col-0, *ROP6-5, prl1-10* and *prl1-10ROP6-5* at 8^th^ day after stratification the *prl1-10ROP6-5* double mutant had elongated narrow, thin and intercalation-lost pavement cell. Scale bar, 50μm. (**I**) Seedlings phenotypes of Col-0, *ROP6-5, prl1-10* and *prl1-10ROP6-5at* 8DAS rose in ½ MS. (**J**) Cotyledon Phenotypes of Col-0 and *prl1-10*. (**K**) Quantitative analysis of cotyledon PCs lobe density of Col-0, *ROP6-5, prl1-10* and *prl1-10ROP6-5* at 4DAS. Data are mean lobe number per μm±SD (n>100 cells from three individual cotyledons). The *prl1* mutant possessed significantly lower density of lobes than Col-0, and *prl1ROP6* had even significantly less lobes density than *prl1* mutant (t test, p<0.001). (**L**) Quantitative analysis of cotyledon PCs neck width of Col-0, *ROP6-5, prl1-10* and *prl1-10ROP6-5* at 4DAS. Data are mean neck width μm±SD (n>100 cells from three individual cotyledons). The *prl1* mutant had significantly wider neck width than Col-0 and also the neck width of *prl1ROP6* was significantly wider than the *prl1* mutant (t test, p<0.001).

Map-based cloning revealed a transition of the 1042^th^ base C to T in the 13^th^ exon of *PRL1* (Pleiotropic Regulatory Locus1, at4g15900), causing its premature translation termination in *L171* (Figure S1 A). The *PRL1* gene encodes a conserved WD40-repeat protein and was reported to respond to hormones, sugar and stress, and function in seedling development and plant immunity (Jia et al., 2017; Li et al., 2018; Monaghan et al., 2009; Nemeth et al., 1998; Palma et al., 2007). To investigate the biological function of *PRL1* in our system, we crossed *L171* to WT to remove the *ROP6-5* background, and the resulting mutant was named *prl1-10*. The homozygous *prl1-10* mutation was confirmed by dCAPs-based analysis with specific primers that are listed in the table S1 (Neff et al., 1998; Neff et al., 2002). The mean pavement cell lobe number and neck width of *prl-10* mutant were 0.67±0.04 and 21.6±0.96μm respectively at 4^th^ day after stratification (Figure 1C, 1K and 1L). The mean lobe number of *prl1-10* mutant decreased significantly (t-test, p<0.001) and the average neck width increased significantly (t-test, p<0.001) when compared to those of Col-0. The reduction of *prl1-10* lobe density shows that *PRL1* plays an important role in the regulation of pavement cell morphogenesis. Interestingly, the lobe numbers were further decreased and the neck width was further increased in *prl1-10 ROP6-5*(*L171*) at 4^th^ day after stratification. However strong overexpression of ROP6 or RIC1 (ROP6 effector) drastically reduced neck widths at later stage of PC development such as 8 DAS (Fu et al., 2005; Fu et al., 2009). These results hint a complex functional interaction of PRL1 with the ROP6 signaling pathway.

From the same ROP6 OX enhancer screen, we isolated another *prl1* allele *prl1-11*, which was also a point mutation causing premature translation termination (mutated at site TGG609TGA of its CDS sequence). In addition, we obtained a T-DNA insertion line *prl1-12* from ABRC (Figure S1A). RT-PCR results showed that expression levels of *PRL1* in these three allelic mutants were severely reduced (Figure S1B). These three allelic *prl1* mutants share similar phenotypes of pavement cell and seedlings (Figure S1C and Figure S1D), indicating that *PRL1* mutation caused decreased pavement cell polarity and *PRL1* could promote pavement cell jigsaw-puzzle formation.

### The *PRL1* mutation enhanced *ROP6* effect on pavement cell morphogenesis

The progression of cell morphogenesis in pavement cells can be divided into three stages. The first and second stages are related to *ROP2*-dependent lateral lobe initiation and growth, the third stage is characterized to be *ROP6*-dependent lateral inhibition of cell outgrowth and longitudinal promotion of cell elongation (Fu et al., 2002; Fu et al., 2009). The puzzle shape of WT pavement cell is regularly formed at 4DAS when *prl1* gets cell contour like what the WT at stage I. When extending time to 8DAS, we found *prl1* PC took the puzzle shape indicating mutation in *PRL1* delayed cell polarization, and the contour of *prl1 ROP6-5* PC was very analogous to those of RIC1 or ROP6 overexpression lines with smooth, long cell outline except for wider cell neck. This result indicated that *prl1* mutation could augment the effect of ROP6 on shaping cell contours at the third stage of cell morphogenesis. The *prl1* was also reported to get enhanced hypersensitivities to variable cues in terms of their developmental phenotypes (Nemeth et al., 1998). So, we predicted that the enhanced hypersensitivities of *prl1* to developmental signals would have similar cellular or molecular mechanisms like its PC to ROP6 signaling.

### PRL1 synergistically interacts with *ROP2* and *ROP6* in regulation of PC morphogenesis

To further explore the genetic relationship between *PRL1* and *ROP6*, we crossed *prl1-10* to *rop6-2* (salk-091737), a *ROP6* loss-of-function mutant, and obtained *prl1-10 rop6-2* double mutant. The mean neck width of *prl1-10 rop6-2* pavement cells was 24.11±0.52μm, greater than that of *rop6-2* (21.4±0.22 μm) and that of *prl1-10* PCs (21.6±0.97μm). Compared with WT (over 100 PCs), the mean lobe number per area in *rop6-2* PCs and *prl1-10* PCs (over 100 PCs) was decreased (Figure 2A, 2G, 2L, and 2M), whereas the lobe density in the *prl1-10 rop6-2* double mutant was greatly decreased compared to both of the single mutants. These genetic analyses suggested *prl-10* and *rop6-2* have synergistic effects on morphogenesis of pavement cells. Whereas apparently the seedling phenotypes of these mutants indicated *prl1-10* was epistatic to *rop6-2* (Figure S3). As described above, however, we also found that *prl1-10* and *ROP6-5* had synergistic effects on the reduction of lobe density and that prl*1-10 ROP6-5 (L171*) possessed wider necks than *prl1-10*, although *ROP6-5* PCs had slightly narrower necks than did WT (Figure 2B, 2C, 2F, 2G, 2H, 2L and 2M). Taken together these results imply that PRL1 and ROP6 may act complicatedly with each other.

**Figure 2.**
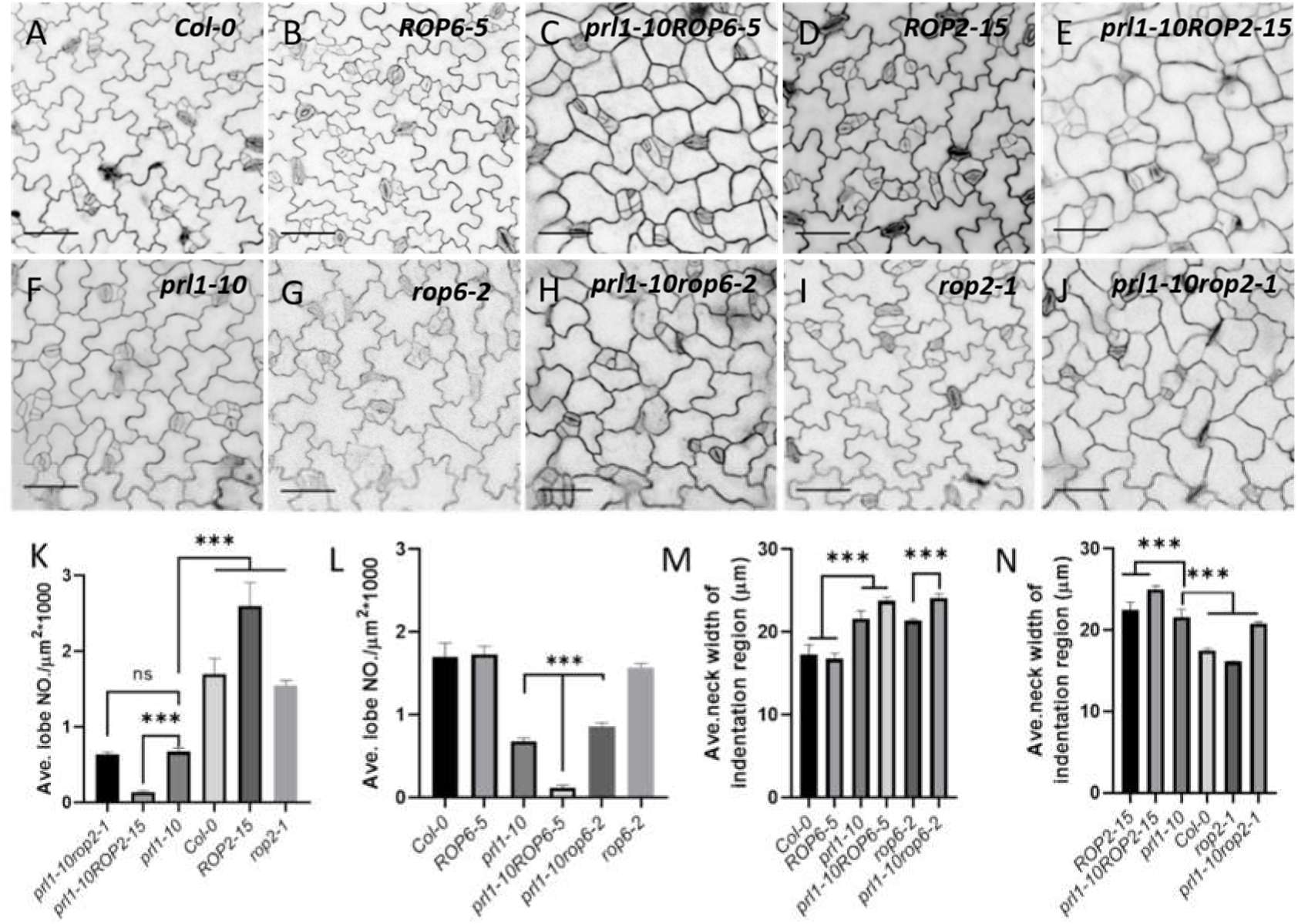
The *prl1* interacts with *ROP2* and *ROP6* synergistically in regulating cellular interdigitation. (**A-J**) The pavement cell contours of variable genetic backgrounds. (**K**) Quantitation of the lobe densities of pavement cell in variable expression levels of ROP2. The lobe density of *prl1rop2* is equal to *prl1* but dramatically less than Col-0 and *rop2*, the mean lobe density of *prl1ROP2* is significantly less than Col-0, *prl1* and *ROP2* (t test, p<0.001, n>100 cells from three individual plants). (**L**) Quantitation of the lobe densities of pavement cell in variable expression levels of *ROP6*. The mean lobe density of *prl1rop6* is higher than *prl1* but significantly less than Col-0 and *rop2*, the lobe density of *prl1ROP6* is significantly less than Col-0, *prl1* and *ROP6* (t test, p<0.001, n>100 cells from three individual plants). (**M**) Quantitation of the neck widths of pavement cells in variable expression levels of *ROP6*. The mean neck width of *prl1rop6* and *prl1ROP6* are significantly wider than *prl1* (t test, p<0.001, n>100 cells from three individual plants). (**N**) Quantitation of the neck widths of pavement cells in variable expression levels of *ROP2*. The mean neck width of *prl1rop2* is less than *prl1*but larger than Col-0 and *rop2-1*, while the neck width of *prl1ROP2* is significantly larger than Col-0, *prl1* and *ROP2* (t test, p<0.001, n>100 cells from three individual plants).

Since *ROP2* and *ROP6* were shown to interact antagonistically in regulation of cell morphogenesis (Fu et al., 2005), we tested whether *ROP2* interacts with *PRL1* in regulating PC morphogenesis. To that end, we constructed *prl1-10 ROP2-15 (GFP-ROP2*) and *prl1-10 rop2-1 (rop2-1*, salk_055328) double mutants. The overexpression of *ROP2* (in the *ROP2-15* line) greatly increased lobe density (Fig 2D, 2M) with 2.60±0.23/1000μm^2^ vs 17.83±0.29μm in WT (Figure 2A, 2M. The mean lobe density of *prl1-10 ROP2-15* PCs was 0.16±0.04/1000μm^2^, which was significantly lower than 0.68±0.04/1000 μm^2^ of *prl1-10* (t test, p<0.001) and *ROP2-15* (Figure 2D, 2E, 2F, 2K and 2N), These results suggest that *prl1-10* and *ROP2* overexpression have synergistic effects on reducing cell polarity in PCs. Similarly, the mean neck width of *prl1-10 ROP2-15* PCs was 25.02±0.44μm, which was wider than 22.45±0.97μm (t test, p<0.001) of *ROP2-15* and 21.06±0.97μm (t test, p<0.001) of *prl1-10* (Figure 2K, 2N). The whole cell contours of *prl1-10 ROP2-15* were similar to those of *CAPOP2-1* except for smaller cell size within *prl1-10ROP2-15* (Figure S2), further suggesting that the *prl1-10* mutation enhanced the effect of ROP2 on PC morphogenesis. Similar to previous observations (Fu et al., 2005), the *rop2-1* knockout mutant exhibited decreased neck width and slightly reduced lobe density in leaf PCs (Figure 2I, 2K and 2N). The neck width and lobe density of *prl1-10 rop2-1* similar to those of *prl1-10* mutant, indicating *PRL1* is epistatic to *ROP2* which was also supported by the phenotypes of their seedlings (Figure S3) and RO2 is necessary for PC morphogenesis of the *prl1*.

Taken together, our results suggest that *PRL1* and ROP2/6 interact synergistically in regulating the jigsaw-puzzle pattern formation of leaf pavement cells.

### The *prl1* mutations lead to differential alteration of ROP2 and ROP6 activities

To further understand the functional relationship between PRL1 and ROP2/6, we examined the activities of ROP2/ROP6 in *prl1-10* mutant. The homozygous *prl1-10 ROP2-15* and *prl1-10 ROP6-5* were utilized for ROP2 and ROP6 activity assay as described previously (Xu et al., 2010). We found that ROP2 activity was increased by *prl1-10*, suggesting PRL1 negatively regulates ROP2 activation (Figure 3A, 3C). Quantitative analysis of active ROP2 indicated that there was 1.5 times more active ROP2 in *prl1-10 ROP2-15* than in *ROP2-15* mutant. We found that ROP6 activity was depressed in *prl1* mutants in contrast to ROP2 activity (Figure 3A, 3C). Active ROP6 decreased by 75% in *prl1-10 ROP6-5* when compared to that in *ROP6-5* (Figure 3B, 3D). The opposing effects of PRL1 on ROP2 and ROP6 activities explain why *prl1-10 rop2-1* double mutants exhibit a similar phenotype that is more or less like *prl1-10* (Fig 2), because ROP2/ROP4 and ROP6 have been confirmed to be antagonistic in PC morphogenesis (Fu et al., 2005; Fu et al., 2009).

**Figure 3.**
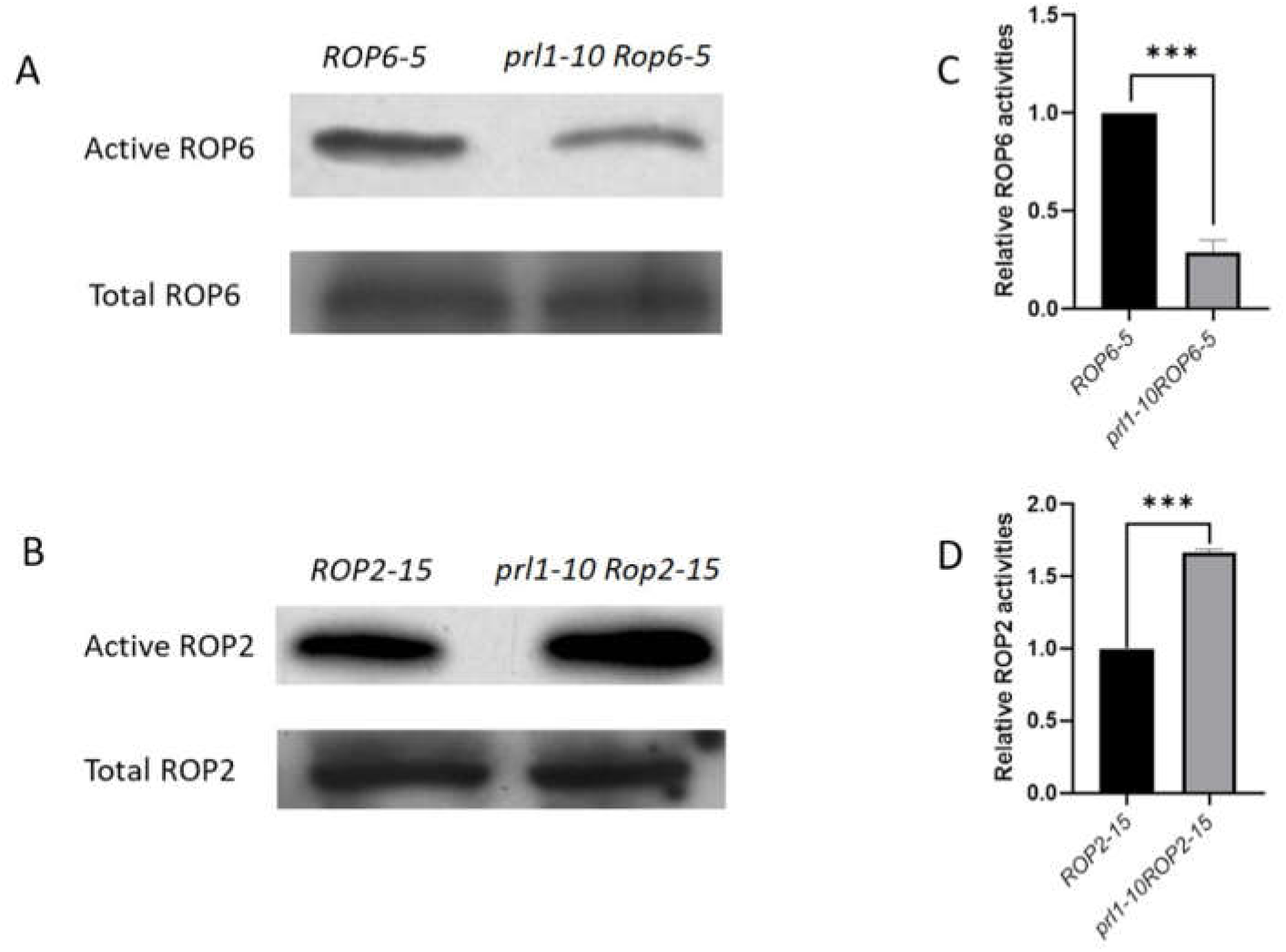
The *PRL1* mutation leads to differential changes of active ROP2 and ROP6. (**A**) Active ROP6 concentrations in the genetic background of *Col-0* and *prl1* mutant. (**B**) Active ROP2 levels in the genetic background of *Col-0* and *prl1* mutant. (**C-D**) Quantification of activities of ROP6 and ROP2 in Col-0 and *prl1* mutant. Data are mean relative concentration of active ROP6 or ROP2 and were standardized to “1” of theses in Col-0.

Taken together, it is concluded that ROP2 and ROP6 are differentially regulated in *prl1*.

### Mutation in *PRL1* specifically causes de-polymerization of actin filaments leading to reduction of PC polarity

To reveal the underlying cellular mechanism for synergistic relationship between *prl1*and *ROP2/ROP6*, we imagined that the basic and essential cytoskeleton organization should have been altered in *prl1* mutant. We crossed cytoskeleton marker lines *GFP-fABD2* and *UBQ::GFP-MBD1* to *prl1-10* respectively and got homozygous *prl1-10 GFP-fABD2 (prl1 GFP-fABD2*) and *prl1-10 UBQ::GFP-MBD1* double mutants. We utilized confocal microscopy to observe structures of cortical MF and MT in pavement cells and found that in the background of *prl1* mutant, in comparison to *GFP-fABD2* and *UBQ::GFP-MBD1*, the cortical actin filaments tended to disassembly while cortical microtubules kept consistent organization and arrays (Figure 4A–4F). So, mutation in *PRL1* could disrupt actin filament other than organization of MT, indicating PRL1 functions in maintaining actin cytoskeleton integrity. Actin filaments in the cortical region of *prl1* cells take diverse depolymerized structures such as continuous but curved filaments, fragmented short filament and even cracked dot-like structures and some accumulated non-filament actin structures (Figure 5M–5P).

**Figure 4.**
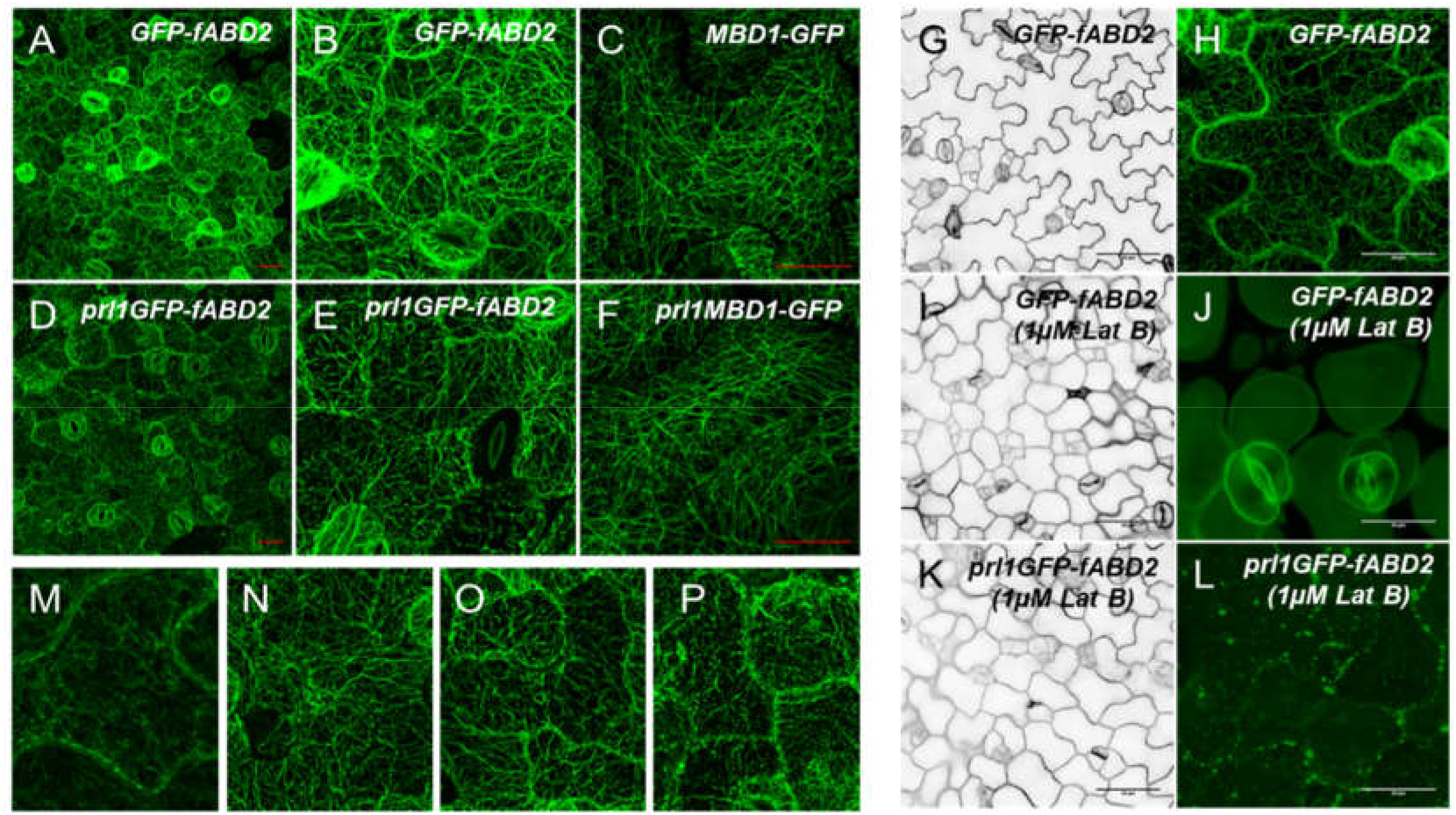
Mutation in PRL1 specifically caused de-polymerization of actin filaments and resultant PC polarity reduction. (**A, B**) Configuration of cortical actin filament in Col-0 with intact and continuous filaments, **B** is the local room-in view of microfilaments in A. Scale bar in **A** is 20 μm. (**D-E, M-P**) Configuration of cortical actin filaments in *prl1* with curved, fragmented and dotted actin structures which were presented in detail in **M**, **N**, **O** and **P**, **E** is the local room-in view of microfilaments in D. Scale bar in D is 20 μm. (**C, F**) Structures of cortical microtubule in Col-0 and *prl1* mutant, there is no difference in cortical microtubule organization between Col-0 and *prl1* mutant. Scale bar, 20 μm. (**G, I, K**) LatB treatment causes Col-0 with *prl1* mutant-like pavement cell phenotype. Scale bar, 50μm. (**H, J, L**) LatB treatment cause completely depolymerization of actin microfilament in Col-0, while in *prl1* depolymerized actin accumulated into punctate structures in cortical region of pavement cell. Scale bar, 20 μm.

**Figure 5.**
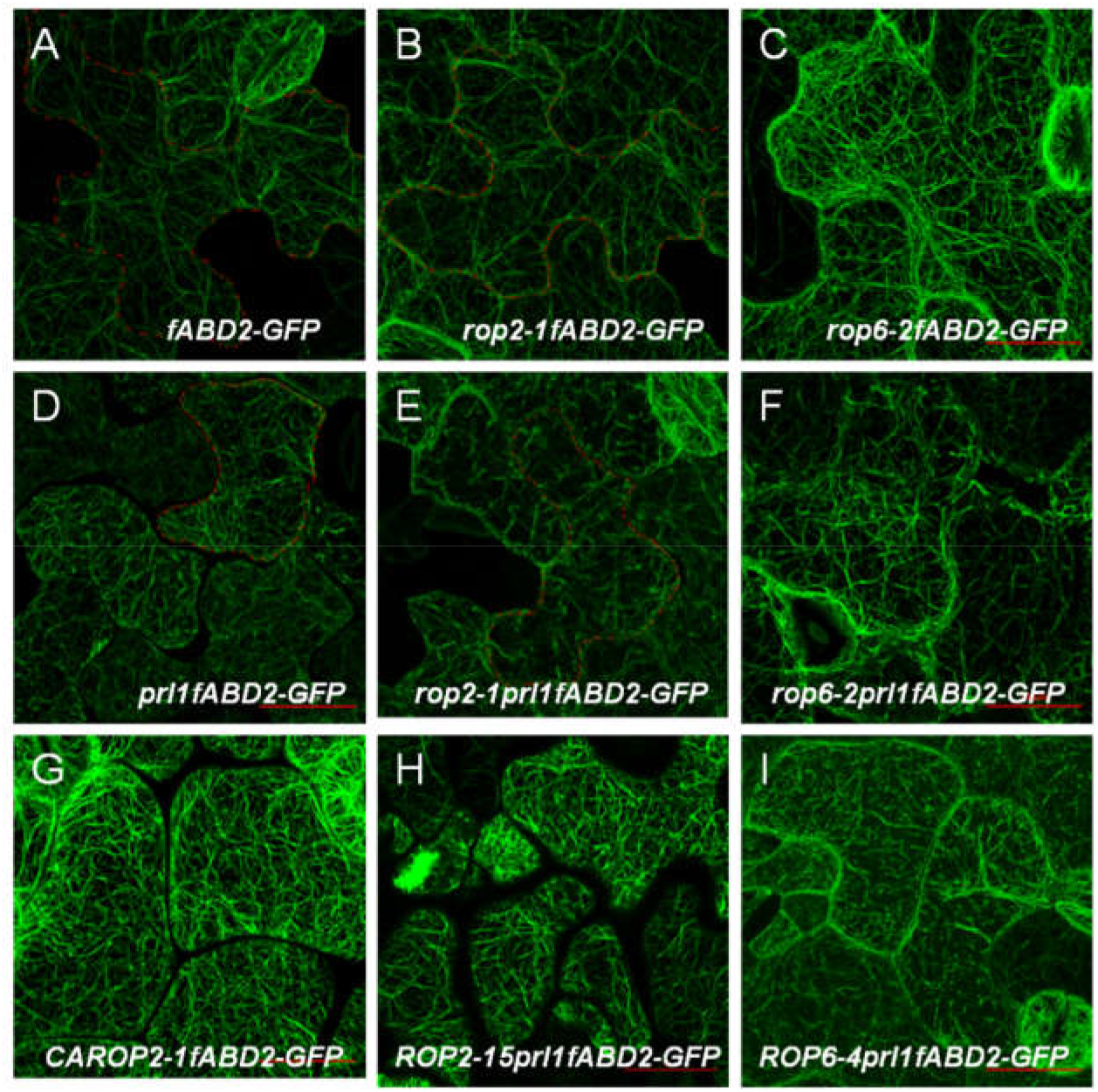
Disassembly of actin filaments in *prl1* mutant is independent of ROP2/6 signaling cascades. (**A**) Cortical region distributed actin filament. (**B**) Cortical actin filaments distribution in the background of rop2-1. (**C**) Cortical actin filaments structures in the background of rop6-2. (**D**) Cortical actin configuration in the background of PRL1 mutation. (**E**) Cortical actin configuration in *rop2-1prl1-10* double mutant. (**F**) Cortical actin configuration in *rop6-2prl1-10* double mutant. (**G**) Cortical actin filament configuration in *CArop2-1*. (**H**) Cortical actin configuration in *ROP2-15prl1-10* double mutant. (**I**) Cortical actin configuration in *ROP6-5prl1-10* double mutant. Scale bar, 50μm.

Could de-polymerized actin filaments be sufficient for the reduction of pavement cell polarity in *prl1* mutant? Next, we used pharmacological method to treat WT seedlings. We germinated *GFP-fADB2* seeds in 1/2MS medium then transferred germinated seeds 1DAS (day after stratification) to LatB-containing 1/2MS medium (1μM) to avoid inhibition of germination by LatB and observed the actin filament structures and pavement cell contours using confocal laser scanning microscopy on 4DAS. The cortical actin filament of treated seedlings completely disassembled and the pavement cell almost lost their lobes analog to *prl1* mutants (Figure 4G–4J). We also observed similar results in LatB-treated *prl1* GFP-*fABD2* mutant.

Unexpectedly, depolymerized actin in *prl1 GFP-fABD2* was stacked and assembled when compared to treated *GFP-fABD2* mutant (Figures 4K–4L).

These results were sufficient to support that *PRL1* mutation causes de-polymerization of actin microfilament and resultantly reduces pavement cell polarity and suggested PRL1 functions in maintaining actin cytoskeleton integrity.

### Actin filaments disassembly in *prl1* mutant is independent of ROP2/6 signaling cascades

Next, we constructed homozygous *rop2-1 GFP-fABD2 and rop6-2 GFP-fABD2* double mutants and *rop2-1 prl1 GFP-fABD2*, *rop6-2 prl1 GFP-fABD2* triple mutants to verify whether ROP2 and ROP6 are involved in actin disassembly in PRL1 signaling. We found cortical actin in cells of *GFP-fABD2* to be continuous, long and smooth filaments (Figure 5A, 5B). The cortical actin in cells of *rop2-1 prl1 GFP-fABD2 and rop6-2 prl1 GFP-fABD2* were short, raptured fragments which were similar to those in *prl1GFP-fABD2* (Figure 5D, 5E and 5F). Those results demonstrated that ROP2 *and ROP6* were not involved in *prl1* caused cortical actin filament disassembly.

What was the mechanism underlying synergistic interaction between *prl1* and ROP2/6? It was proposed that the two genes functioning separately but finally converging at a node of one given biological progress would be interact synergistically (Perez-Perez et al., 2009). Thus, we constructed homologous *ROP2-15 prl1 GFP-fABD2* and *ROP6-5 prl1 GFP-fABD2* triple mutants for observation of MF. By means of confocal laser scanning microscopy, we observed that there was more fragmented actin which was evenly distributed across the cortical region in comparison to *prl1 GFP-fABD2* (Figure 5H, 5I). These results indicated that *ROP2* and *ROP6* were probably involved in not only reorganizing monomer actin into filamentous configuration but also facilitating accumulation actin in the cellular cortical region. In early stage of cell morphogenesis, *CAROP2-1*, *RIC1*, *prl1 ROP2* and *prl1 ROP6* possess similar regular cell phenotypes, which hints the probability of their similar functions in promotion of actin rearrangement and accumulation in cortical cell region (Figure 2C, 2E, 5G and S2). So, we argued that PRL1 negatively governed two separate signaling pathways, one is ROP-independent actin disassembly and the other is ROP2/6-dependent actin polymerization, finally converged in synergistic reorganization actin configuration and distribution.

### The *prl1* mutation mimics stresses-controlled activation of ROP2/6

What signals directs differential change of activities of ROP2 and ROP6 in *prl1* mutant? It was reported that null mutant in *prl1* could de-repress the expression of stress-related genes (Salchert et al., 1998), indicating *prl1* mutation initiates stress signals. We hypothesized that *prl1* mutation-arisen stress would be the initial signal relaying to activation of ROP2 and ROP6. To test this hypothesis, we conducted followed experiments. First, we germinated *prl1-10* in ½ MS for 1 day, then transferred to 150nM NaCl-containing ½ MS and rose for 3days. Next, we observed pavement cell contours at 4DAS. We found that contours of chloride sodium-treated *prl1-10* pavement cells are identical to those of *CArop2-1* at 4 DAS except for smaller cell size (Figure 6A – 6D), indicating stress could change cell polarity. Next, we tested active ROP2 and ROP6 in seedlings grown in 150mM chloride sodium-containing ½ MS medium and found similar activity changes of ROP2 and ROP6 to *prl1* (Figure 6E and 6F). We also found similar changes of ROP2 and ROP6 in darkness (Figure 6E and 6F). In addition, we found that actin was assembled into bundles and actin bundle levels were proportional to stress degrees (Figure 6G). We also observed evenly distributed thicker but fragmented actin bundles over cortical region in pavement cell of *prl1 mutant* which were raised in 150 mM chloride sodium-containing ½ MS medium (Figure 6G). These results supported that stress could successively change ROP2/6 signaling cascades-dependent cortical actin organization. In combination with previous data, in *prl1* mutant differential changes of activities of ROP2/6 would be the result of *prl1* mutation-initiated stresses.

**Figure 6.**
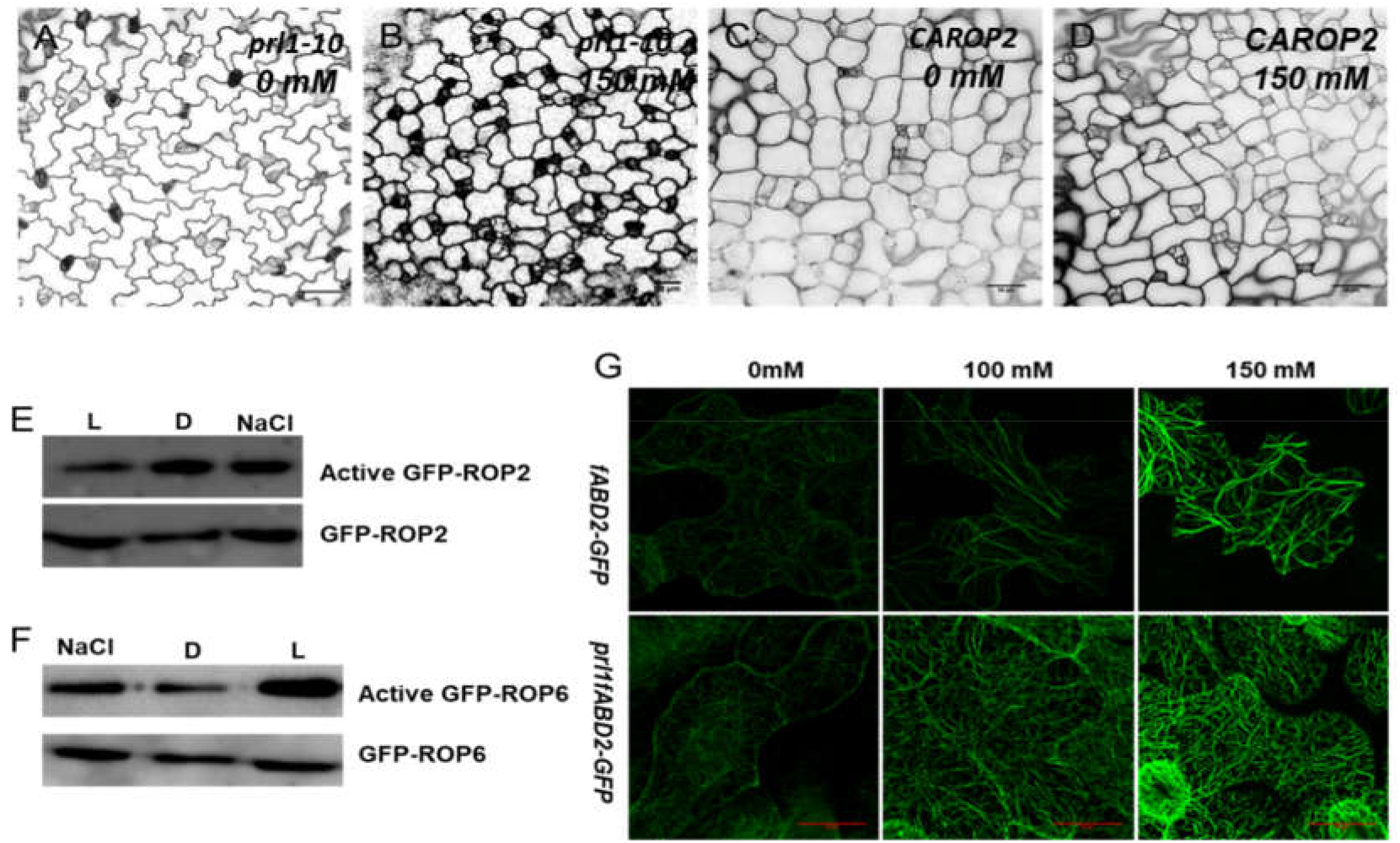
The *prl1* mutation-arisen stress answers for differential activation of ROP2/6. (**A-D**) Pavement cell phenotype of *prl1* and *CArop2-1* mutants. In B and D, seeds were germinated for one day in incubation chamber, and then were transferred to 150 mM NaCl-containing ½ MS and cultured for 3 days before observation. Scale bar, 50μm. (**E-F**) Stresses could change activities of ROP2 and ROP6 as in *prl1* mutant. (**G**) Actin structures in Col-0 and *prl1* background treated with different concentration of NaCl (mM). The actin bundling degree is dependent on levels of stress intensity. Scale bar, 50 μm.

## Discussion

### *PRL1* negatively-governed *ROP*-dependent *and* -independent signaling pathways interact synergistically in regulating actin cytoskeleton organization for cell morphogenesis

Two types of actin binding proteins (ABPs) including polymerization factors and de-polymerization factors govern separately and collaboratively successive actin dynamic process in response to upstream various signals. A bulk of data has been provided to define these two types of proteins in organizing actin filament, for example formin, profilin and ADF/Cofilin had been mentioned to function synergistically to affect rapid turnover of actin filaments (Michelot et al., 2007), but what signals were passed on to them to guide ABPs-dependent actin filament dynamic was scarcely described. In this study, we found PRL1 negatively controlled two parallel signaling pathways to maintain dynamic actin homeostasis at normal physiological condition (Figure 7), although we do not know exactly the mechanical mechanism of PRL1 maintaining actin integrity yet. We did get that mutation in *PRL1* broke the steady state of cortical actin filament turnover with preference to be de-polymerized. Depolymerized actin would be competent to be reorganized into higher order actin configurations in response to upstream signals (Burke et al., 2014; Suarez and Kovar, 2016). In fact, it was found *prl1* mutant was hypersensitive to various cues (Nemeth et al., 1998), what the depolymerized actin might be the cellular mechanism of these hypersensitivities in responses to variable cues. Additionally, we found moderate change in activity of ROP6 or ROP2 in background of *prl1* could lead to obvious changes of pavement cell contours by promoting isotopic cell outgrowth, which were largely based on actin reorganization. In consistent to Fu’s conclusion in role of actin for cell outgrowth (Fu et al., 2002), we get new role of ROP6, like ROP2, in actin filament reorganization and redistribution which in turn promotes lateral cell outgrowth in early the 1st stage of pavement cell development. While at the later stage of pavement cell development, we proposed the hypersensitivity of *prl1* to ROP6 with narrow, long and smooth pavement cell, in large degree, was attribute to indirect effect of depolymerized actin filament on pavement cell morphogenesis by changing interaction between ROP6-directed organization of actin cytoskeleton and tubulin through releasing inhibition of MF on distribution and reconfiguration of MT. Our result still supports previous findings that ROP2 and ROP6 functions antagonistically, even in background of *prl1*, as *rop2-1 prl1-10* took cell contour more like *prl1-10* while *prl1-10 rop6-2* more like *prl1-10 ROP2-5*. Although ROP2 and ROP6 could change cell contours and actin configurations in *prl1* mutant, but they fail in restoring actin into filamentous structures indicating the basic role of PRL1 is maintaining actin integrity while ROP2/ROP6 signaling pathways may be side effect of PRL1 mutation or negative feedback of actin depolymerization.

**Figure 7.**
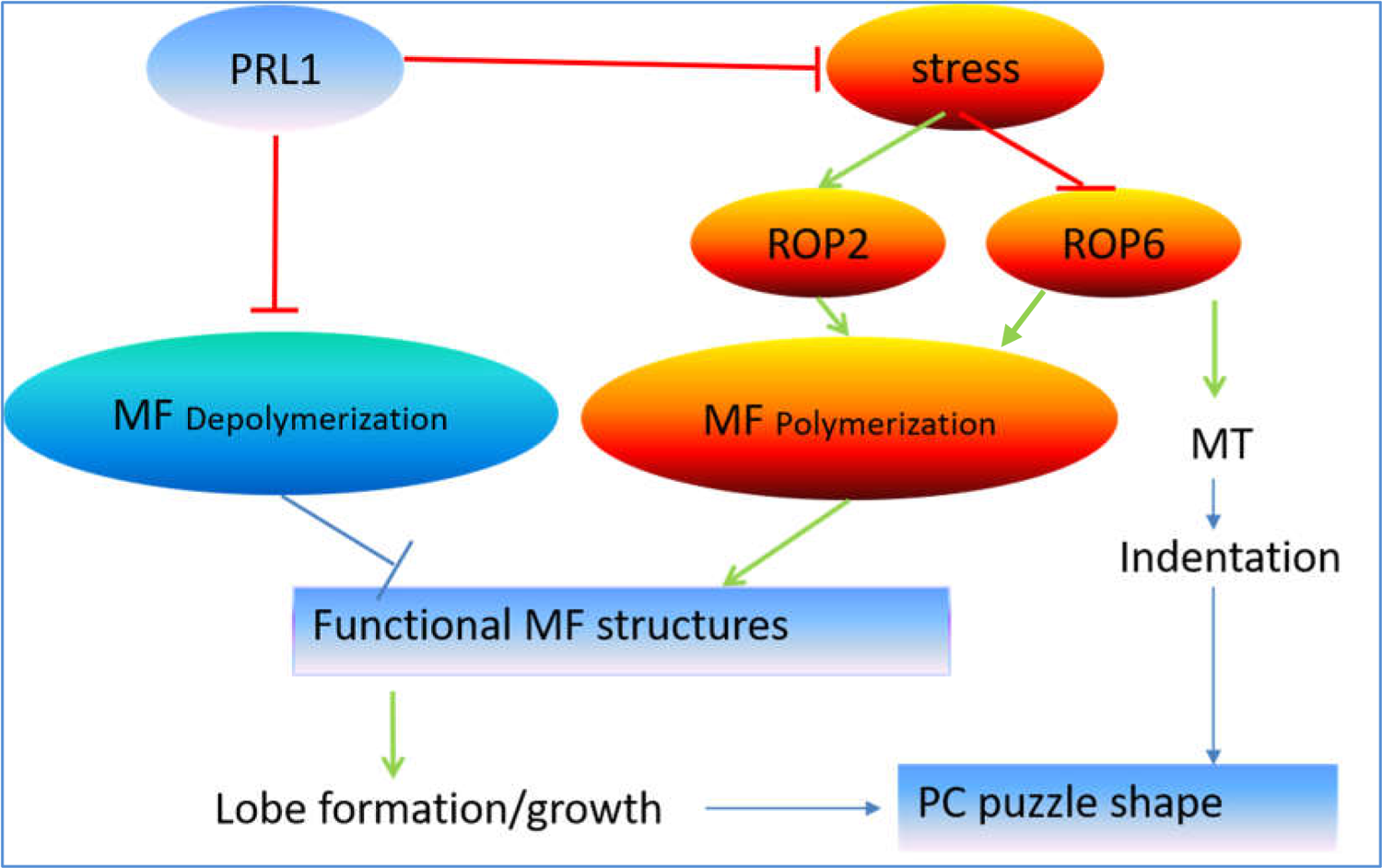
Proposed model for *PRL1* in PC morphogenesis. PRL1 is responsible for dynamic homeostasis of actin filaments. It negatively manages actin depolymerization and controls stress-directed ROPs activities-dependent actin polymerization/bundling. Stresses could differentially regulate activities of ROP2 and ROP6 which are both able to promote actin organization. PRL1 mutation caused actin disassembly is epistatic to ROPs-dependent actin remodeling and finally leads to reduction of polarity of pavement cell.

In considering of stress signals on actin cytoskeleton organization, it had been revealed that plants could be resistant to certain degree of stresses by promoting formation of stress pretreatment-induced actin bundles (Wang et al., 2010). Our results may establish a ROPs-dependent link between stresses and cytoskeleton dynamic. As higher levels of actin organization could make plant more resistant to further stresses, our result makes new knowledge that ROP-dependent actin cytoskeleton assembly would be a kind of cellular mechanisms of plants for further resistance to more intensive stresses after salt acclimation.

To summarize, in the schematic model we present that PRL1 negatively integrates two separate ROP-dependent and -independent signaling pathways in actin dynamic regulation and contributing to polar pavement cell morphogenesis in Arabidopsis.

### ROP2 and ROP6 mark stereotypical microfilament-promoting cell outgrowth

It is well-accepted opinion that cell differentiation and growth are concomitant programs to each other. In early stage, pavement cells take isotropic outgrowth, while in later stage they make for differentiation by directional growth along longitudinal cell axis. PM-localized ROP2 and ROP6 antagonize mutually in two levels for directing cell morphogenesis, one is active ROP2 and ROP6 competing for PM localization and forming discrete PM distribution, the other is negatively regulating actions of their contrary downstream effectors (Fu et al., 2005; Fu et al., 2009). The biological function of ROP2 was early defined in actin filament assembly and locally lateral cell outgrowth, matching the general concept of “cell growth and cell differentiation are concomitant”, while the biological function of ROP6 was delimited to specify cortical cellular microtubule array for local outgrowth inhibition in the later stage of pavement cell morphogenesis. Whereas, the observed pavement cell phenotypes of *ROP6* in the background of *prl1* mutant implied that *ROP6* signaling cascade was also involved in cell outgrowth in early growing stage of cell morphogenesis. In present study, we found an additional biological role of ROP6, like ROP2, in reorganization of actin filament for cell lateral outgrowth in early stage of pavement cell development then transiting to the third stage of ROP6-mediated MT organization and lateral cell growth inhibition. In combination of relationship between *ROP2* and *ROP6*, we propose that ROPs-directed actin filament organization-dependent isotropic cell lateral outgrowth is a canonical program for later cell polar outgrowth. This hypothesis reinforced the basic knowledge that cell growth is functionally prerequisite for polar cell development.

### The general biological significance of discovery of PRL1 Mutation-caused MF depolymerization

*PRL1* was suggested to be a central regulator in many biological functions including root, leaf and flower development, plant innate immunity, carbohydrate metabolism, cell polarity development and even cellular energy homeostasis (Abraham et al., 2003; Flores-Perez et al., 2010; Li et al., 2007; Nemeth et al., 1998; Palma et al., 2007; Salchert et al., 1998). The biochemical and molecular functions of PRL1 were mainly defined in cullin4 (CUL4) E3 ligase as negatively specific receptor for SnRK1 turnover and in NTC (Prp19 complex)/MAC (MOS4-associated complex) for transcription regulation. There are gaps between its molecular roles and physiological functions, although gene transcription and cell energy deprivation were generally thought to be responsible for cell development and innate immunity (Yang et al., 2014). In the present study, the new identified role of PRL1 controlling actin modulation would contribute to our further understanding of the molecular mechanism of PRL1 underlying its physiological roles. Accumulating evidences indicated actin cytoskeleton played important roles in plant innate immunity (Henty-Ridilla et al., 2014; Henty-Ridilla et al., 2013; Moral et al., 2017; Underwood, 2016). Plant evolved two major defense strategies, PTI and ETI against pathogen infection (Day et al., 2011; Kobayashi and Hakuno, 2003; Rajamuthiah and Mylonakis, 2014). Actin organization and rearrangement would be the wrestling target of host cells and pathogens for their respective purposes. First, actin polymerization should be responsible for plant resistance to pathogen infection. For example, LatB inhibited actin polymerization promoted significant increase of *Pseudomonas syringae* infection to host plants (Henty-Ridilla et al., 2013). In response to pathogen infection, plant host cells organize actin filament into higher density of actin filament network over cortical cellular region to physically prevent their invasion or into actin filament trajectory for antimicrobial components transporting to invasion site to defeat pathogen penetration into host cell (Janda et al., 2014; Jelenska et al., 2014; Underwood, 2016). Whereas, during evolution of pathogen-host interaction, pathogens have evolved specific strategies to achieve their infection such as hijacking or mimic actions of the intracellular machinery ABPs to regulate actin dynamics and modulate organization of the actin filament network, such as actin filament depolymerizing factors such as ADFs (Fu et al., 2014; Guan et al., 2013; Henty-Ridilla et al., 2014; Kang et al., 2014; Porter et al., 2012; Shimono et al., 2016; Tian et al., 2009). In *prl1* mutant, actin cytoskeleton exhibited depolymerized status. On the one hand, it decreased the defense ability of host cell; on the other hand, the depolymerized actin became competent to be hijacked by pathogens for their successful infection. Therefore, many studies found hypersensitivity of *prl1* and other core NTC/MAC components mutants to *Pseudomonas syringae and Hyaloperonospora parasitica* (*H.p*.) (Monaghan et al., 2009; Palma et al., 2007). So, actin cytoskeleton organizing role of PRL1 should be the cause for plant innate immunity.

Another well documented function of PRL1 is regulating gene expression by affecting accumulation of microRNA and small interfering RNA (siRNA) and alternative splicing pre-mRNA (Jia et al., 2017; Li et al., 2018; Zhang et al., 2014). Here, new discovered role of PRL1 in maintaining cellular actin cytoskeleton homeostasis could also be the cause of transcriptional alternation. In addition to extensive distribution and function in cytoplasm, actin also localized and performed diverse functions in nuclear including transcription regulation (Bettinger et al., 2004; Chen and Shen, 2007; Hofmann et al., 2004; Virtanen and Vartiainen, 2017). Here, the form of depolymerized actin filaments was consistent with functional form in regulation of gene transcription (Virtanen and Vartiainen, 2017). Further, it was also found that monomeric or disassembled actin could be conveniently transported into nuclear by binding to ADF/cofilin and change transcription (Pendleton et al., 2003). So, actin depolymeriztion could be the cause of transcriptional alterations in *prl1* mutant. Although we did not distinguish cause-effect relationship between actin depolymerization and stress initiation, *prl1*-triggered stress could also be another factor for its transcription alteration and immunity defects.

Taken together, our finding suggested that *PRL1* directed a specific signaling pathway which could affect actin filament dynamic homeostasis and cellular stress status that could be the link between polar cell development, plant cell innate immunity and even gene transcription regulation. In the light of evolutionarily conservation, PRLG1, the homolog of PRL1 in mammal cells, even Prp19 complex /MAC complex should also have the conserved function in keeping dynamic homeostasis of cortical cellular cytoskeleton (Kleinridders et al., 2009). Thereafter, our results in plant would advance understanding of the molecular mechanism underlying conserved PRL1/PLRG1 biological functions.

## Material and Methods

### Plant materials and growth conditions

All plants used in this study are Col-0 ecotype. The *prl1-10* and *prl1-12* are first isolated single nucleotide replacement premature translation termination mutants by ourselves. The *rop2-1* and *rop6-2* are T-DNA insertional mutants salk_091737C and salk_055328C ordered from ABRC. The ROP2-15 (ref.) and ROP6-5 and RIC1-3 OE lines and CArop2 (Fu et al., 2005) were regularly used in ZB Yang’s lab. To observe cytoskeleton configuration, we used the cytoskeleton marker lines *GFP-fABD_2_* (Sheahan et al., 2004) and *UBQ::GFP-MBD* (Eng and Wasteneys, 2014) and crossed them respectively with *prl1-10*, *rop2-1*, *rop6-2*, *ROP2-15*, *ROP6-5* and *CArop2* to construct homozygous double or triple mutants. Plants were grown at 22°C on ½ MS agar plate and later transferred to soil with 16hr-light/8hr-dark cycles unless indicated otherwise. The seedlings were photographed and images were edited with photoshop2018.

### Sodium chloride and low temperature treatment

To observe effect of stress on shape of cotyledon pavement cell, we used 100nm and 150nm sodium chloride-containing ½ MS agar plate to raise plants. After germinated for 1 day on ½ MS agar plate, seedlings were transferred to variable concentration of sodium chloride-containing ½ MS agar plate for further 3 days growth, then the seedlings were used for followed cell shape observation, ROP activity assay and actin cytoskeleton configuration imaging. To evaluate effect of general stress on ROPs activity, we use low temperature (4°C) to treat 4DAS seedlings for 12hr, then extracted total proteins and evaluated ROPs’ activities using pull-down assay.

### Mapping-based location of candidate mutation of PRL1

To localize the causal mutation in genome of *L171* mutant, the *L171* (Col-0 ecotype) was out-crossed to WT of Landsberg *erecta* (Ler) to construct recombinant inbred lines (RIL). As the causal mutation in *L171* was identified to be recessive, in F2 population the leaves of single *prl1*-like plant were collected individually. Genomic DNA was extracted from leaves of each single plant using the CTAB method separately. Pool the DNA samples for bulked segregate analysis. Design and synthesize primers to amplify InDel (Monsanto Arabidopsis polymorphism) sequence-containing DNA segments to confirm genetic linkage between markers and potential mutation. After first-pass mapping, the mutation of interest was positioned to region in the 4^th^ chromosome of Arabidopsis between BAC clones of T6G15 (corresponding to SNP_Name CER460892) and F9F13(CER466358), then synthesize more primers according to InDel markers to perform fine-mapping. The causal mutation was delimited to a region within FCAALL and further narrowed down to regions between FCA3 clone and FCA5 clone. All genes in this region were sought out and compared phenotype of L171 with allelic mutants of these genes, amplified the genomic DNA sequence of suspected gene in L171 and found the causal mutation (the 1042^th^ base C to T in the 13^th^ exon of *PRL1*)which causes premature coding termination of PRL1, additional allelic *prl1* mutant were collected with identical phenotypes.

### Confocal microscopy analysis of Pavement cell contour in cotyledon

Since propidium iodide (PI) can be taken up and stain plant cell wall, we use PI staining to present cell shape. We cut the 4-DAS cotyledons of plants grown at 22°C on ½ MS agar plate and immerged them in 10μg/ml PI solution for 5-10 Mins, then observed and imaged pavement cell shape with confocal microscopy (Zeiss LSM 880). PI was excited using 561nm laser light and the emitted florescent light was collected between 600nm and 655nm.The images were edited by ImageJ.

### Pull-down assay of ROP’s activity

To analyze activities of ROP2 and ROP6, we used pull-down in combination with western blot method. As RIC1 is commonly specific target of activated ROP2 (GTP-bound) and ROP6 (GTP-bound), we use excessive MBP-RIC1 to affinity ROP2 and ROP6 separately and quantify relative amount of activated ROP2 to total ROP2 (including GDP-bound and GTP-bound) and GTP-ROP6 to total ROP6 (including GDP-bound and GTP-bound) using western blot. Total protein was extracted from 1 gram 8 DAS seedlings (without and with physiological treatment) grown on ½ MS plate, including *GFP-ROP2* (*ROP2-15*), GFP-ROP6 (ROP6-5), *prl1 ROP2-15* and *prl1 ROP6-5* seedlings. Twenty microgram of MBP-RIC-conjugated agarose beads were added to the total protein extracts, and incubated 3hrs or overnight at 4°C. The beads were washed 3 times at 4 °C (5min/time), GTP-bound GFP-ROP2 or -ROP6 that was associated with the MBP-RIC1 beads was used for analysis by western blotting with anti-GFP antibody (Santa Cruz Biotechnology, Santa Cruz, CA). Meanwhile, a fraction of total protein was analyzed by immunoblot to quantify total GFP-ROP2 or -ROP6 (including GDP-bound and GTP-bound). The amount of GTP-bound ROP2 or ROP6 was normalized to that of total ROP2 or ROP6 respectively. The level of GTP-bound ROPs in *prl1* or physiological treatment plants relative to controls (ROP2-5 or ROP6-5) was calculated by dividing the amount of normalized GTP-bound ROP2 or ROP6 from *prl1* genetic background lines or each treatment by the normalized amount from the control, which is normalized to be “1”.

### Visualization of Actin and Microtubule with Confocal Microscopy

To visualize the cortical actin configuration and microtubule configuration, we used laserscanning confocal microscopy LSM 880 with Airyscan software to image the cotyledon pavement cells of variable genetic background plants with identical parameters. 4-DAS plant cotyledon were cut and mounted on the slide, added distilled water and covered with coverslip and observed under Zeiss LSM 880. We collected Z stacks of optical sections and generated Z-series maximum intensity projections of GFP fluorescence to evaluate actin and microtubule configurations.

## Acknowledgements

Thank Dr. Wenfeng, Sun for technique support on pull-down assay. We are grateful to Prof.& Dr. Shuang, Wu for his stimulating discussion and critical comments on this manuscript. Thank funding for X.G. by NSF from Fujian province (2017J01599), and for Z.Y. by start-up fund from FAFU and for Y.F. by the Major Research Plan of the National Natural Science Foundation of China (90817105).

## Author contributions

X.G, Y.F and Z.Y. conceived and designed the study. X.G., B.Y, J.Z., C.W. and X.W. performed the experiments and made data analysis. X.G. wrote the manuscript. All authors read and approved the final version of the manuscript.

## Supplementary materials

**Figure S1.**
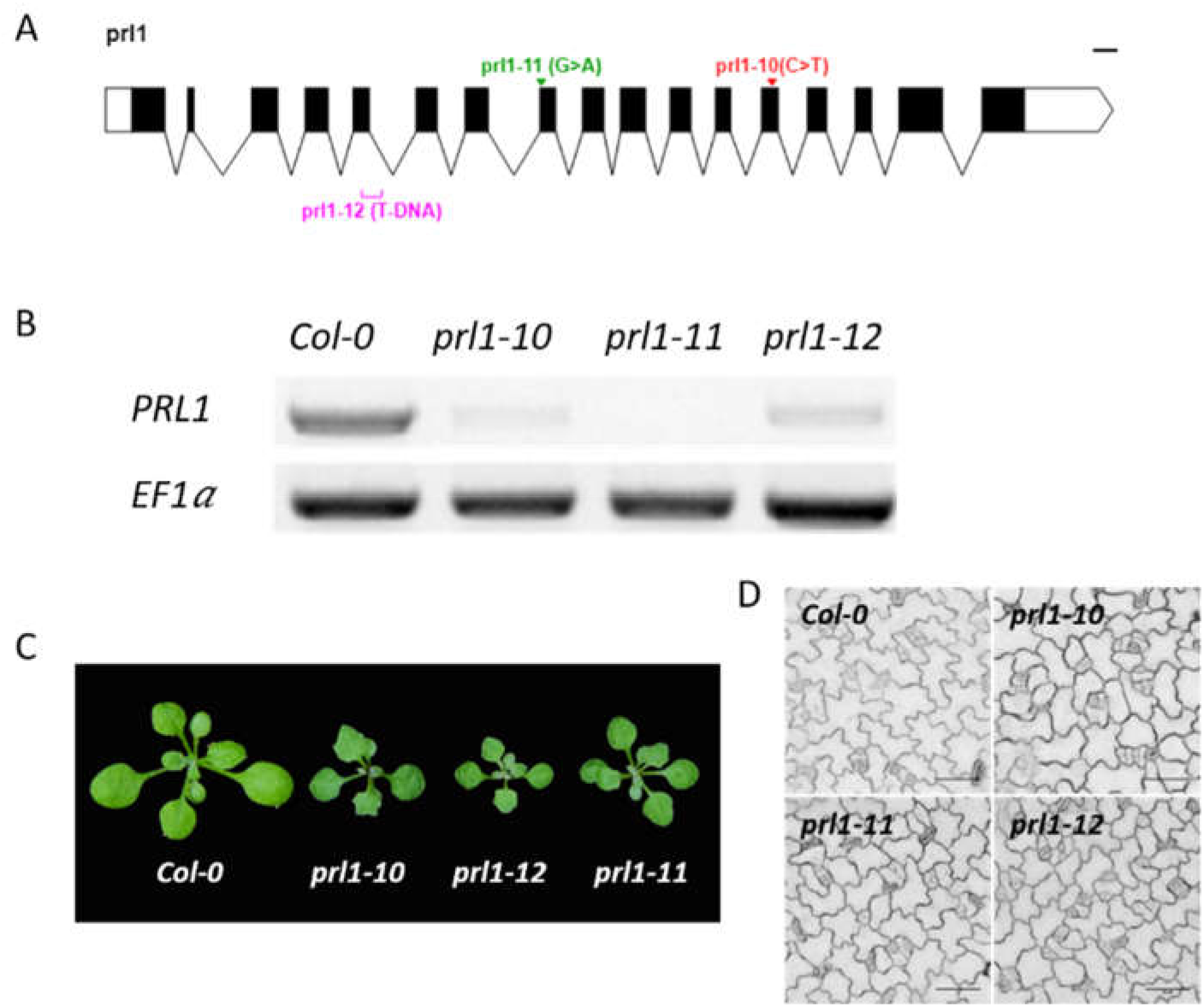
Allelic PRL1 mutants share similar phenotypes. (A) Diagram of structures of *prl1* allelic genes. The bar on right top of the diagram indicates 100bp. (B) Expression levels of *PRL1* gene in WT and three different *prl1* allele, EF1a gene as control. (C) 14-DAS seedlings of WT, *prl1-10, prl1-11* and *prl1-12*. (D) 4-DAS pavement cell contours in WT, *prl1-10, prl1-11* and *prl1-12*.

**Figure S2.**
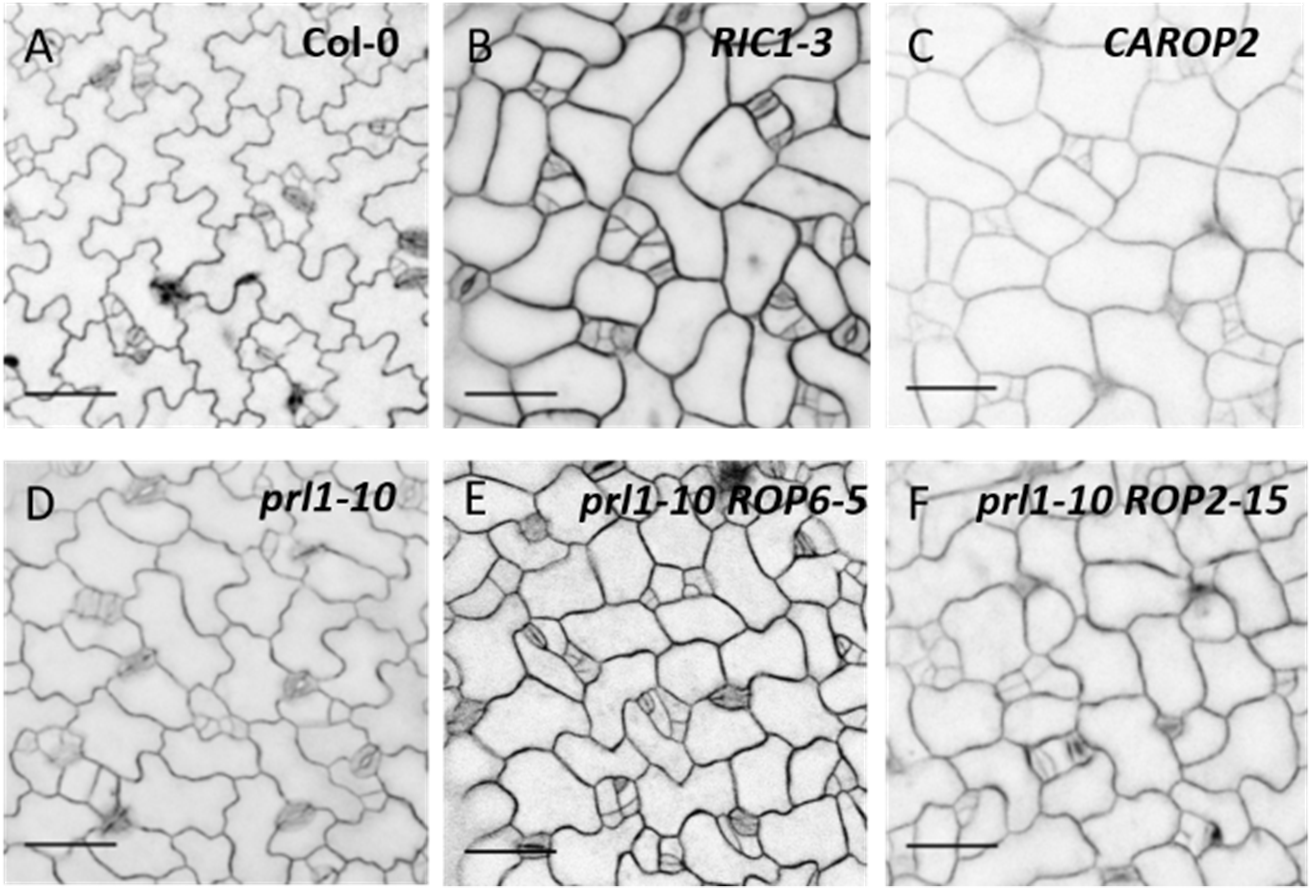
Pavement cell shapes with variable genetic background. (A) 4-DAS pavement cell of WT. (B) 4-DAS pavement cell of *RIC-OE* line *RIC1-3*. (C) 4-DAS pavement cell of *ROP2* constitutive activation line. (D) 4-DAS pavement cell of *prl1-10*. (E) 4-DAS pavement cell of homologous *prl1-10 ROP6-5* double mutant. (F) 4-DAS pavement cell of homologous *prl1-10 ROP2-15* double mutant.

**Figure S3.**
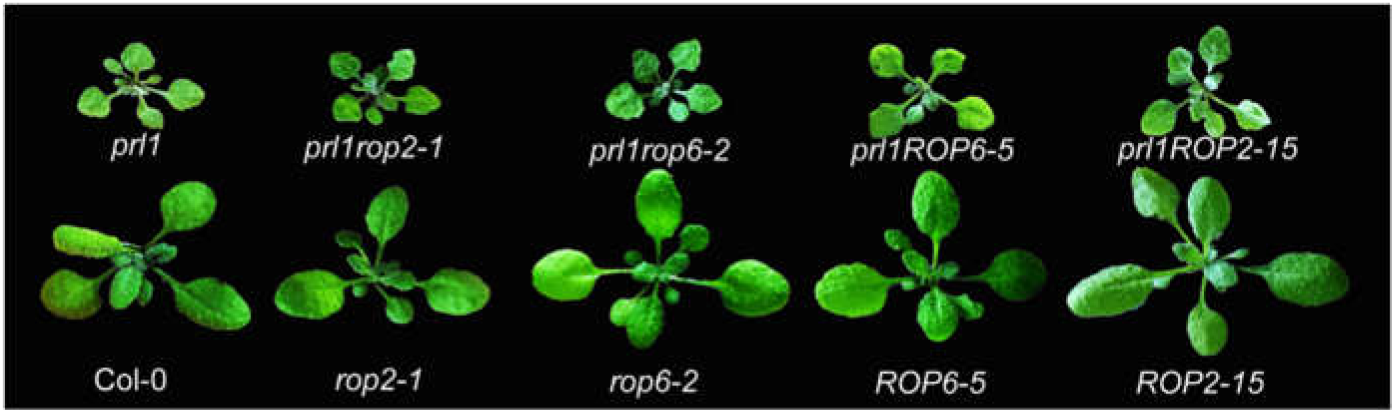
PRL1 is epistasis to ROPs on development of Arabidopsis.

**Table S1.**
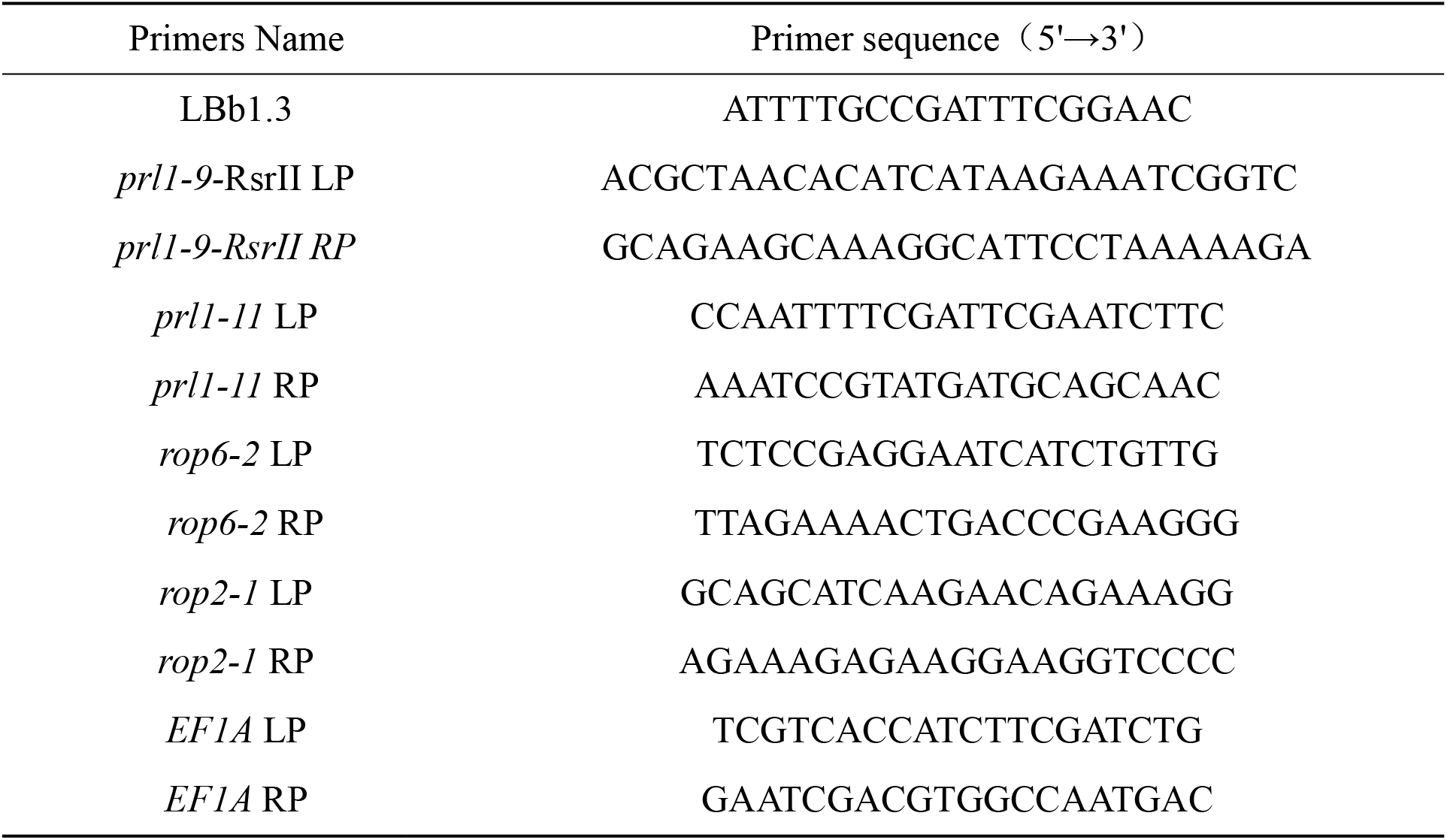
Primers used for genotype identification

## Reference

Abraham, E., Rigo, G., Szekely, G., Nagy, R., Koncz, C., and Szabados, L. (2003). Light-dependent induction of proline biosynthesis by abscisic acid and salt stress is inhibited by brassinosteroid in Arabidopsis. Plant molecular biology 51, 363–372.

Andrianantoandro, E., and Pollard, T.D. (2006). Mechanism of actin filament turnover by severing and nucleation at different concentrations of ADF/cofilin. Molecular cell 24, 13–23.

Baruah, A., Simkova, K., Hincha, D.K., Apel, K., and Laloi, C. (2009). Modulation of O-mediated retrograde signaling by the PLEIOTROPIC RESPONSE LOCUS 1 (PRL1) protein, a central integrator of stress and energy signaling. The Plant journal: for cell and molecular biology 60, 22–32.

Bettinger, B.T., Gilbert, D.M., and Amberg, D.C. (2004). Actin up in the nucleus. Nature reviews Molecular cell biology 5, 410–415.

Burke, T.A., Christensen, J.R., Barone, E., Suarez, C., Sirotkin, V., and Kovar, D.R. (2014). Homeostatic actin cytoskeleton networks are regulated by assembly factor competition for monomers. Current biology: CB 24, 579–585.

Campellone, K.G., and Welch, M.D. (2010). A nucleator arms race: cellular control of actin assembly. Nature reviews Molecular cell biology 11, 237–251.

Chang, M., Li, Z., and Huang, S. (2017). Monomeric G-actin is uniformly distributed in pollen tubes and is rapidly redistributed via cytoplasmic streaming during pollen tube growth. The Plant journal: for cell and molecular biology 92, 509–519.

Chhabra, E.S., and Higgs, H.N. (2007). The many faces of actin: matching assembly factors with cellular structures. Nature cell biology 9, 1110–1121.

Edwards, M., Zwolak, A., Schafer, D.A., Sept, D., Dominguez, R., and Cooper, J.A. (2014). Capping protein regulators fine-tune actin assembly dynamics. Nature reviews Molecular cell biology 15, 677–689.

Chen, M., and Shen, X. (2007). Nuclear actin and actin-related proteins in chromatin dynamics. Current opinion in cell biology 19, 326–330.

Day, B., Henty, J.L., Porter, K.J., and Staiger, C.J. (2011). The pathogen-actin connection: a platform for defense signaling in plants. Annual review of phytopathology 49, 483–506.

Etienne-Manneville, S., and Hall, A. (2002). Rho GTPases in cell biology. Nature 420, 629–635.

Eng, R.C., and Wasteneys, G.O. (2014). The microtubule plus-end tracking protein ARMADILLO-REPEAT KINESIN1 promotes microtubule catastrophe in Arabidopsis. The Plant cell 26, 3372–3386.

Flores-Perez, U., Perez-Gil, J., Closa, M., Wright, L.P., Botella-Pavia, P., Phillips, M.A., Ferrer, A., Gershenzon, J., and Rodriguez-Concepcion, M. (2010). Pleiotropic regulatory locus 1 (PRL1) integrates the regulation of sugar responses with isoprenoid metabolism in Arabidopsis. Molecular plant 3, 101–112.

Fu, Y., Duan, X., Tang, C., Li, X., Voegele, R.T., Wang, X., Wei, G., and Kang, Z. (2014). TaADF7, an actin-depolymerizing factor, contributes to wheat resistance against Puccinia striiformis f. sp. tritici. The Plant journal: for cell and molecular biology 78, 16–30.

Fu, Y., Gu, Y., Zheng, Z., Wasteneys, G., and Yang, Z. (2005). Arabidopsis interdigitating cell growth requires two antagonistic pathways with opposing action on cell morphogenesis. Cell 120, 687–700.

Fu, Y., Li, H., and Yang, Z. (2002). The ROP2 GTPase controls the formation of cortical fine F-actin and the early phase of directional cell expansion during Arabidopsis organogenesis. The Plant cell 14, 777–794.

Fu, Y., Xu, T., Zhu, L., Wen, M., and Yang, Z. (2009). A ROP GTPase signaling pathway controls cortical microtubule ordering and cell expansion in Arabidopsis. Current biology: CB 19, 1827–1832.

Guan, X., Buchholz, G., and Nick, P. (2013). The cytoskeleton is disrupted by the bacterial effector HrpZ, but not by the bacterial PAMP flg22, in tobacco BY-2 cells. Journal of experimental botany 64, 1805–1816.

Henty-Ridilla, J.L., Li, J., Day, B., and Staiger, C.J. (2014). ACTIN DEPOLYMERIZING FACTOR4 regulates actin dynamics during innate immune signaling in Arabidopsis. The Plant cell 26, 340–352.

Henty-Ridilla, J.L., Shimono, M., Li, J., Chang, J.H., Day, B., and Staiger, C.J. (2013). The plant actin cytoskeleton responds to signals from microbe-associated molecular patterns. PLoS pathogens 9, e1003290.

Higaki, T., Akita, K., and Hasezawa, S. (2020). Elevated CO2 promotes satellite stomata production in young cotyledons of Arabidopsis thaliana. Genes to cells: devoted to molecular & cellular mechanisms 25, 475–482.

Hofmann, W.A., Stojiljkovic, L., Fuchsova, B., Vargas, G.M., Mavrommatis, E., Philimonenko, V., Kysela, K., Goodrich, J.A., Lessard, J.L., Hope, T.J., et al. (2004). Actin is part of preinitiation complexes and is necessary for transcription by RNA polymerase II. Nature cell biology 6, 1094–1101.

Huang, S., Qu, X., and Zhang, R. (2015). Plant villins: versatile actin regulatory proteins. Journal of integrative plant biology 57, 40–49.

Humphries, J.A., Vejlupkova, Z., Luo, A., Meeley, R.B., Sylvester, A.W., Fowler, J.E., and Smith, L.G. (2011). ROP GTPases act with the receptor-like protein PAN1 to polarize asymmetric cell division in maize. The Plant cell 23, 2273–2284.

Hussey, P.J., Ketelaar, T., and Deeks, M.J. (2006). Control of the actin cytoskeleton in plant cell growth. Annual review of plant biology 57, 109–125.

Janda, M., Matouskova, J., Burketova, L., and Valentova, O. (2014). Interconnection between actin cytoskeleton and plant defense signaling. Plant signaling & behavior 9, e976486.

Jelenska, J., Kang, Y., and Greenberg, J.T. (2014). Plant pathogenic bacteria target the actin microfilament network involved in the trafficking of disease defense components. Bioarchitecture 4, 149–153.

Jia, T., Zhang, B., You, C., Zhang, Y., Zeng, L., Li, S., Johnson, K.C.M., Yu, B., Li, X., and Chen, X. (2017). The Arabidopsis MOS4-Associated Complex Promotes MicroRNA Biogenesis and Precursor Messenger RNA Splicing. The Plant cell 29, 2626–2643.

Kang, Y., Jelenska, J., Cecchini, N.M., Li, Y., Lee, M.W., Kovar, D.R., and Greenberg, J.T. (2014). HopW1 from Pseudomonas syringae disrupts the actin cytoskeleton to promote virulence in Arabidopsis. PLoS pathogens 10, e1004232.

Kleinridders, A., Pogoda, H.M., Irlenbusch, S., Smyth, N., Koncz, C., Hammerschmidt, M., and Bruning, J.C. (2009). PLRG1 is an essential regulator of cell proliferation and apoptosis during vertebrate development and tissue homeostasis. Mol Cell Biol 29, 3173–3185.

Kobayashi, I., and Hakuno, H. (2003). Actin-related defense mechanism to reject penetration attempt by a non-pathogen is maintained in tobacco BY-2 cells. Planta 217, 340–345.

Kudryashova, E., Heisler, D.B., and Kudryashov, D.S. (2017). Pathogenic Mechanisms of Actin Cross-Linking Toxins: Peeling Away the Layers. Current topics in microbiology and immunology 399, 87–112.

Li, H., Xu, T., Lin, D., Wen, M., Xie, M., Duclercq, J., Bielach, A., Kim, J., Reddy, G.V., Zuo, J., et al. (2013). Cytokinin signaling regulates pavement cell morphogenesis in Arabidopsis. Cell research 23, 290–299.

Li, J., Blanchoin, L., and Staiger, C.J. (2015). Signaling to actin stochastic dynamics. Annual review of plant biology 66, 415–440.

Li, J., and Staiger, C.J. (2018). Understanding Cytoskeletal Dynamics During the Plant Immune Response. Annual review of phytopathology 56, 513–533.

Li, R., and Gundersen, G.G. (2008). Beyond polymer polarity: how the cytoskeleton builds a polarized cell. Nature reviews Molecular cell biology 9, 860–873.

Li, S., Liu, K., Zhou, B., Li, M., Zhang, S., Zeng, L., Zhang, C., and Yu, B. (2018). MAC3A and MAC3B, Two Core Subunits of the MOS4-Associated Complex, Positively Influence miRNA Biogenesis. The Plant cell 30, 481–494.

Li, Y., Deng, M., Liu, H., Li, Y., Chen, Y., Jia, M., Xue, H., Shao, J., Zhao, J., Qi, Y., et al. (2020). ABNORMAL SHOOT 6 interacts with KATANIN 1 and SHADE AVOIDANCE 4 to promote cortical microtubule severing and ordering in Arabidopsis. Journal of integrative plant biology.

Li, Y., Smith, C., Corke, F., Zheng, L., Merali, Z., Ryden, P., Derbyshire, P., Waldron, K., and Bevan, M.W. (2007). Signaling from an altered cell wall to the nucleus mediates sugar-responsive growth and development in Arabidopsis thaliana. The Plant cell 19, 2500–2515.

Liu, X., Yang, Q., Wang, Y., Wang, L., Fu, Y., and Wang, X. (2018). Brassinosteroids regulate pavement cell growth by mediating BIN2-induced microtubule stabilization. Journal of experimental botany 69, 1037–1049.

Pollard, T.D. (2007). Regulation of actin filament assembly by Arp2/3 complex and formins. Annual review of biophysics and biomolecular structure 36, 451–477.

Michelot, A., Berro, J., Guerin, C., Boujemaa-Paterski, R., Staiger, C.J., Martiel, J.L., and Blanchoin, L. (2007). Actin-filament stochastic dynamics mediated by ADF/cofilin. Current biology: CB 17, 825–833.

Monaghan, J., Xu, F., Gao, M., Zhao, Q., Palma, K., Long, C., Chen, S., Zhang, Y., and Li, X. (2009). Two Prp19-like U-box proteins in the MOS4-associated complex play redundant roles in plant innate immunity. PLoS pathogens 5, e1000526.

Moral, J., Montilla-Bascon, G., Canales, F.J., Rubiales, D., and Prats, E. (2017). Cytoskeleton reorganization/disorganization is a key feature of induced inaccessibility for defence to successive pathogen attacks. Molecular plant pathology 18, 662–671.

Neff, M.M., Neff, J.D., Chory, J., and Pepper, A.E. (1998). dCAPS, a simple technique for the genetic analysis of single nucleotide polymorphisms: experimental applications in Arabidopsis thaliana genetics. The Plant journal: for cell and molecular biology 14, 387–392.

Neff, M.M., Turk, E., and Kalishman, M. (2002). Web-based primer design for single nucleotide polymorphism analysis. Trends in genetics: TIG 18, 613–615.

Nemeth, K., Salchert, K., Putnoky, P., Bhalerao, R., Koncz-Kalman, Z., Stankovic-Stangeland, B., Bako, L., Mathur, J., Okresz, L., Stabel, S., et al. (1998). Pleiotropic control of glucose and hormone responses by PRL1, a nuclear WD protein, in Arabidopsis. Genes & development 12, 3059–3073.

Palma, K., Zhao, Q., Cheng, Y.T., Bi, D., Monaghan, J., Cheng, W., Zhang, Y., and Li, X. (2007). Regulation of plant innate immunity by three proteins in a complex conserved across the plant and animal kingdoms. Genes & development 21, 1484–1493.

Pendleton, A., Pope, B., Weeds, A., and Koffer, A. (2003). Latrunculin B or ATP depletion induces cofilin-dependent translocation of actin into nuclei of mast cells. The Journal of biological chemistry 278, 14394–14400.

Perez-Perez, J.M., Candela, H., and Micol, J.L. (2009). Understanding synergy in genetic interactions. Trends in genetics: TIG 25, 368–376.

Porter, K., Shimono, M., Tian, M., and Day, B. (2012). Arabidopsis Actin-Depolymerizing Factor-4 links pathogen perception, defense activation and transcription to cytoskeletal dynamics. PLoS pathogens 8, e1003006.

Rajamuthiah, R., and Mylonakis, E. (2014). Effector triggered immunity. Virulence 5, 697–702.

Salchert, K., Bhalerao, R., Koncz-Kalman, Z., and Koncz, C. (1998). Control of cell elongation and stress responses by steroid hormones and carbon catabolic repression in plants. Philosophical transactions of the Royal Society of London Series B, Biological sciences 353, 1517–1520.

Sheahan, M.B., Staiger, C.J., Rose, R.J., and McCurdy, D.W. (2004). A green fluorescent protein fusion to actin-binding domain 2 of Arabidopsis fimbrin highlights new features of a dynamic actin cytoskeleton in live plant cells. Plant physiology 136, 3968–3978.

Shimono, M., Lu, Y.J., Porter, K., Kvitko, B.H., Henty-Ridilla, J., Creason, A., He, S.Y., Chang, J.H., Staiger, C.J., and Day, B. (2016). The Pseudomonas syringae Type III Effector HopG1 Induces Actin Remodeling to Promote Symptom Development and Susceptibility during Infection. Plant physiology 171, 2239–2255.

Sit, S.T., and Manser, E. (2011). Rho GTPases and their role in organizing the actin cytoskeleton. Journal of cell science 124, 679–683.

Smith, L.G., and Oppenheimer, D.G. (2005). Spatial control of cell expansion by the plant cytoskeleton. Annual review of cell and developmental biology 21, 271–295.

Staiger, C.J., and Blanchoin, L. (2006). Actin dynamics: old friends with new stories. Current opinion in plant biology 9, 554–562.

Staiger, C.J., Sheahan, M.B., Khurana, P., Wang, X., McCurdy, D.W., and Blanchoin, L. (2009). Actin filament dynamics are dominated by rapid growth and severing activity in the Arabidopsis cortical array. The Journal of cell biology 184, 269–280.

Suarez, C., and Kovar, D.R. (2016). Internetwork competition for monomers governs actin cytoskeleton organization. Nature reviews Molecular cell biology 17, 799–810.

Szymanski, D., and Staiger, C.J. (2018). The Actin Cytoskeleton: Functional Arrays for Cytoplasmic Organization and Cell Shape Control. Plant physiology 176, 106–118.

Tian, M., Chaudhry, F., Ruzicka, D.R., Meagher, R.B., Staiger, C.J., and Day, B. (2009). Arabidopsis actin-depolymerizing factor AtADF4 mediates defense signal transduction triggered by the Pseudomonas syringae effector AvrPphB. Plant physiology 150, 815–824.

Underwood, W. (2016). Contributions of host cellular trafficking and organization to the outcomes of plant-pathogen interactions. Seminars in cell & developmental biology 56, 163–173.

Van Troys, M., Huyck, L., Leyman, S., Dhaese, S., Vandekerkhove, J., and Ampe, C. (2008). Ins and outs of ADF/cofilin activity and regulation. European journal of cell biology 87, 649–667.

Virtanen, J.A., and Vartiainen, M.K. (2017). Diverse functions for different forms of nuclear actin. Current opinion in cell biology 46, 33–38.

Wang, C., Zhang, L., Yuan, M., Ge, Y., Liu, Y., Fan, J., Ruan, Y., Cui, Z., Tong, S., and Zhang, S. (2010). The microfilament cytoskeleton plays a vital role in salt and osmotic stress tolerance in Arabidopsis. Plant biology 12, 70–78.

Xia, G., Ramachandran, S., Hong, Y., Chan, Y.S., Simanis, V., and Chua, N.H. (1996). Identification of plant cytoskeletal, cell cycle-related and polarity-related proteins using Schizosaccharomyces pombe. The Plant journal: for cell and molecular biology 10, 761–769.

Xu, T., Wen, M., Nagawa, S., Fu, Y., Chen, J.G., Wu, M.J., Perrot-Rechenmann, C., Friml, J., Jones, A.M., and Yang, Z. (2010). Cell surface-and rho GTPase-based auxin signaling controls cellular interdigitation in Arabidopsis. Cell 143, 99–110.

Yang, S., Tang, F., and Zhu, H. (2014). Alternative splicing in plant immunity. International journal of molecular sciences 15, 10424–10445.

Yang, Z. (2008). Cell polarity signaling in Arabidopsis. Annual review of cell and developmental biology 24, 551–575.

Zhang, S., Liu, Y., and Yu, B. (2014). PRL1, an RNA-binding protein, positively regulates the accumulation of miRNAs and siRNAs in Arabidopsis. PLoS genetics 10, e1004841.

Zhou, Z., Shi, H., Chen, B., Zhang, R., Huang, S., and Fu, Y. (2015). Arabidopsis RIC1 Severs Actin Filaments at the Apex to Regulate Pollen Tube Growth. The Plant cell 27, 1140–1161.

